# Meta-analysis of Genes and Pathways that Protect Against Hypoxia

**DOI:** 10.64898/2026.08.05.743086

**Authors:** Emmett McGranaghan, Ginger Z. Watzinger, Norton Kerri-Ann, Dana L. Miller, Heather L. Bennett

**Affiliations:** Trinity College, Department of Biology, 300 Summit Street, Hartford, CT, United States of America; Computational Sciences Program, Reem-Kayden Center for Science and Computation, Bard College, New York. United States of America; Department of Biochemistry, University of Washington, School of Medicine, Seattle, Washington, United States of America

**Keywords:** Hypoxia, Anoxia, Mice, *C. elegans*, Yes Associated Protein, yap-1, *mxl-3*, MAX (MYC associated transcriptional regulator X) *ador-1*, adenosine receptor 1

## Abstract

Oxygen is essential for all terrestrial animals, but there is dramatic variability in how well different animals and even different cell types can adapt to reduced oxygen availability. We used a meta-analysis of the literature, with a focus on mouse studies, to identify pathways that might act to protect animals in low oxygen environments. We identified 108 genes whose mRNA levels change under hypoxia, and 55 genes critical for mounting a response to hypoxia. With this data, we developed a list of conserved genes, and we tested three *C.elegans* genes previously uncharacterized in hypoxia, *mxl-3*, *yap-1,* and *ador-1*, and found that loss of function altered egg-laying during and after hypoxia. Our method provides a more targeted approach of how to screen for hypoxic phenotypes and study in more genetically tractable organisms to show mechanisms.

## Introduction

Oxygen is essential for all known terrestrial animals. Oxygen is the final electron acceptor in oxidative phosphorylation, a major source of cellular adenosine triphosphate (ATP), and is required for the function of hundreds of other cellular enzymes. Reduced oxygen delivery to tissues, hypoxic ischemia, is a hallmark of many diseases and health conditions, including stroke, myocardial infarction, and pulmonary embolism (Menger and Vollmar 2007). But not all cells and tissues are equally sensitive to hypoxic ischemia ( Hochachka et al. 1996; Michiels 2004; Nystul and Roth; 2004; Storey and Storey 2004), suggesting that there are mechanisms that can protect against the damaging effects of hypoxia.

When animals are confronted with decreased oxygen availability (hypoxia), they often adapt to shift from a reliance on oxidative phosphorylation toward glycolysis to ensure continued production of ATP. One key factor that directs the response to hypoxia is the conserved hypoxia-inducible transcription factor, HIF-1 (Powell-Coffman et al. 1998; Jiang et al. 2001; Semenza 2001; Shen et al. 2005; Kaelin and Ratcliffe 2008)((Powell-Coffman 2010). Orthologues of HIF1 have been found in all metazoans to date (Loenarz et al. 2011). When oxygen becomes limiting, activation of HIF-1 leads to increased production of glycolytic proteins, to support production of ATP, as well as various stress response pathways to mitigate the accumulation of cellular damage. However, the genes that are regulated by HIF1 in hypoxia vary in different cells and organisms (Storey and Storey 2004), and there are some responses to hypoxia that are independent of HIF-1 (Bishop et al. 2004; Shen et al. 2006; Budde and Roth 2010). For example, *C. elegans*, *Drosophila*, and zebrafish embryos all enter into a reversible state of suspended animation, where all observable biological activity arrests, but (at least in *C. elegans*) suspended animation doesn’t require HIF1 (Foe and Alberts 1985; Padilla and Roth 2001; Padilla et al. 2002; Nystul and Roth 2004).

*C. elegans* is a particularly powerful model for delineating the molecular factors that mediate adaptive responses to hypoxia. Because they are small, *C. elegans* do not rely on a circulatory system to deliver oxygen to cells and tissues; instead, cells are in direct contact with the gaseous environment (Dusenbery 1980; Jiang et al. 2001; Shen et al. 2005). This allows precise experimental control of the concentration of oxygen experienced by cells. Studies in *C. elegans* show that the specific behavioral and physiological responses to hypoxia depend on how much oxygen is in the environment (Iranon and Miller 2012). When placed in a gradient, *C. elegans* prefers environments with ∼10% oxygen (Dusenbery 1980; Voorhies and Ward 2000) room air is ∼21% O_2_ (de Bono and Bargmann 1998; de Bono et al. 2002; Gray et al. 2004). In moderate hypoxia, which we defined as 0.5%-3.5% O_2_, metabolic activity is limited by the concentration of O_2_ and HIF-1 is required for embryo survival and for continued postembryonic development (Voorhies and Ward 2000; Nystul and Roth 2004; Miller and Roth 2009). Suspended animation is triggered in anoxic environments (operationally defined as < 10 ppm O_2_;(Padilla et al. 2002). In embryos, suspended animation requires the conserved spindle assembly checkpoint protein SAN-1 (Nystul et al. 2003). SAN-1 is not required for embryos to survive moderate hypoxia (0.5% O_2_), which is lethal for *hif-1* mutant embryos (Padilla et al. 2002; Nystul et al. 2003; Hajeri et al. 2005; Hajeri et al. 2008). Similarly, *hif-1* is not required for embryos to survive suspended animation (Nystul et al. 2003).

Our goal is to more fully delineate the molecular genetic features of conserved adaptive responses to hypoxia. We used a meta-analysis of the published literature to identify genes that have been shown to play a role in the response to hypoxic ischemia in mammalian models. We identified three candidate genes for further testing based on this analysis: *mxl-3, a* Myc-associated transcription factor, *ador-*1, an adenosine G-protein coupled receptor, and *yap-1,* a Yes-associated transcription factor. We assayed how mutants in each of these genes responded to different hypoxic conditions. We found that all three candidates were able to survive and recover from anoxia-induced suspended animation. However, we did observe a defect in *mxl-3,* and *yap-1* mutant animals exposed to moderate hypoxia; when exposed to 0.5% O_2_, these mutants were not able to maintain reproductive capacity. We show that we can identify mammalian genes with roles in hypoxia, and study in genetically tractable organisms providing insights into mechanisms of regulation.

## Experimental Methods

### Literature Search

We surveyed the literature in “PubMed” and “Google Scholar”, from the last 25 years and identified studies of hypoxic conditions where altered gene function, either loss of gene function, gain of gene function, or changes in gene expression led to a survival advantage to oxygen deprivation. We use the keywords in no specific combination “hypoxia,” “anoxia,” “ischemia” “conservation,” “mouse studies,” “oxygen deprivation,” “rodents studies,” “adaptation,” and “survival advantage” and “protection” to identify genes that are involved in protection from hypoxia. The literature review included, but was not limited to, candidate gene reported, genome-wide association studies, transcriptional profiling, microarray analysis reported to alter survival to hypoxia and ischemia in mouse samples, *M. musculus.* To ensure that we compiled an extensive list of hypoxia protective genes, at least two independent co-authors searched the literature using this approach blinded to previously identified candidate genes and literature databases.

A database was assembled in Microsoft Excel with the NCBI Gene ID, gene name across species (human homolog, as well as *C. elegans* ortholog), *in vitro* or *in* vivo experimental analysis for each gene identified, impact of gene function on hypoxia survival. We included data from studies conducted as 1) functional tests where researchers tested a target gene or specific genes and described its effect on survival to hypoxia or 2) transcriptomics studies where hypoxic conditions were created and researchers performed whole organism or single cell transcriptomic changes to identify broad gene expression changes. All data used in the analysis are included in the manuscript as supplemental files or uploaded to the online journal repository.

### Human Homolog and *Caenorhabditis elegans* Ortholog Identification

Homologs and orthologs of genes were identified using Genome Alliance and DIOPT were used to confirm human and *C. elegans* homologs and orthologs.

### Reactome Cell Pathway Analysis and Gene Ontology Assessment

Cellular pathway analysis was performed for lists of mouse genes using Reactome, which provides details on the broad cross species relationships and molecular interactions determined by human biology. Mouse candidate genes were input into Reactome. A complete list of all the cell pathways identified is provided within the supplement. Gene ontology for enriched biological processes was performed using the online bioinformatics databases GOrilla Enrichment Analyses with a Fisher’s test false discovery rate set to 0.01 followed by a Bonferroni correction test (Eden et al. 2007; Eden et al. 2009). All data can be found in supplement or provided upon request.

### *C. elegans* Maintenance and Strains

Strains were maintained at 20 degrees Celsius on NGM plates seeded with OP50 *E. coli*. Some strains were obtained directly from CGC, others were obtained from the National BioResource Project in Tokyo Japan. The following strains were used in this work: N2, referred to as wild type, *hif-1(ia4)*, *yap-1(tm1416)*, *mxl-3(ok1947)*, *ador-1(yum1524)*, *ador-1(gk961191)* 0x backcrossed.

### Anaerobic Exposure, Assessment of Survival, and Measurement of Egg Laying

*C. elegans* on NGM plates seeded with OP50 were placed into environmental chambers, and gas containing the indicated concentration of O_2_ was perfused continuously for the duration of the exposure as in (Miller and Roth 2009; Fawcett et al. 2012). Palmitic acid (10 mg/mL in ethanol) was used to ring the plates; when the ethanol evaporates it leaves a crunchy barrier of palmitic acid crystals that prevents animals from leaving the surface of the plate. For exposure at L4, animals were picked from mixed populations grown at room temperature for at least one generation. For L1, adults were allowed to lay eggs for 2-6 h and the embryos were allowed to hatch at room temperature for 16-20h before hypoxic exposure. Alternatively, L4 were picked to NGM plates seeded with OP50 and the next morning the plates were flooded with sterile M9 and newly-hatched L1 were recovered by mouth pipet. Wild-type (N2) controls were included with every experiment. Before each experiment, samples were numbered randomly so that the genotype was blinded when data were collected, the samples were not decoded until after all data was collected.

For hypoxic conditions, tanks of compressed O_2_ gasses were mixed by Airgas (Seattle, WA) and certified standard to within 2% of O_2_ content of either 1000ppm O_2_ or 5000ppm O_2_ with the balance N_2_. Pure N_2_ (<10 ppm O_2_) were used for anoxic conditions. Environmental chambers were pyrex crystallization dishes that were drilled and fit with plastic male luer-to-hose barb fittings (Cole Parmer) secured with epoxy for gas entrance and venting. Glass plates were used as lids, sealed with Dow Corning vacuum grease. In some instances, environments were in 2.5L AnaeroPack System Jars (Mitsubishi Gas Chemical Co., Inc), drilled and fit with plastic male luer-to-hose barb fittings (Cole Parmer) secured with epoxy. Flow tubes (number 032-41^st^, Aalborg) were used for constant flow rate and pressure, and gas was delivered through one-eighth-inch outer-diameter FEP or nylon tubing (Cole Parmer) with connection by hose-barb snap-connectors (Cole Parmer) or brass compression fittings (Seattle Fluid Systems). Gas was hydrated with dH_2_O before delivery to chambers using a 125 mL gas wash bottle with a fritted cylinder (VWR).

Animals were exposed to hypoxic or anoxic conditions at room temperature. L4 exposed to hypoxia were scored after recovering in room air for 24-36h to assess survival, developmental progression to normal-looking gravid adults, and for the protruding vulva phenotype. L4 that did not survive had clearly lost turgor pressure and did not respond to mechanical stimulation. Some animals survived but appeared scrawny and unhealthy, but test experiments indicate these animals were generally fertile; the number of animals with the scrawny appearance was noted for all samples. *C. elegans* exposed to hypoxia as L1 were scored after 48-60h for viability to gravid adult. In some experiments, the protruding vulva phenotype was also scored.

### Statistical Analysis and Illustration Design

Statistical significance for Reactome pathway analysis was determined using a binomial test. For all anoxia and hypoxia experiments, statistical analysis was performed using GraphPad Prism version 11 (GraphPad Software, La Jolla, CA). Statistical significance is determined using either a one-way Anova or two-way Anova with a Bonferroni or Tukey correction. We report the standard deviation of the mean for all experiments. Illustrations designed using BioRender.

## Results

To identify genes and cellular pathways that regulate and protect against the deleterious effects of hypoxic and ischemic insults in mice we first conducted a literature review. We initially focused on the vertebrate model organism mouse; the cellular, molecular, and genetic approaches available allowed for identification of genes and pathways with functional tests to more accurately describe regulation of response. In addition, the advanced technological approaches allowed investigation of more recent studies, *i.e.,* those conducted within the last 25 years. We surveyed the literature PubMed databases at the National Center for Biotechnology Information (NCBI), as well as open-source platform Google Scholar. Candidate genes were identified using a set of targeted keywords, see Figure 1.

**Figure 1:**
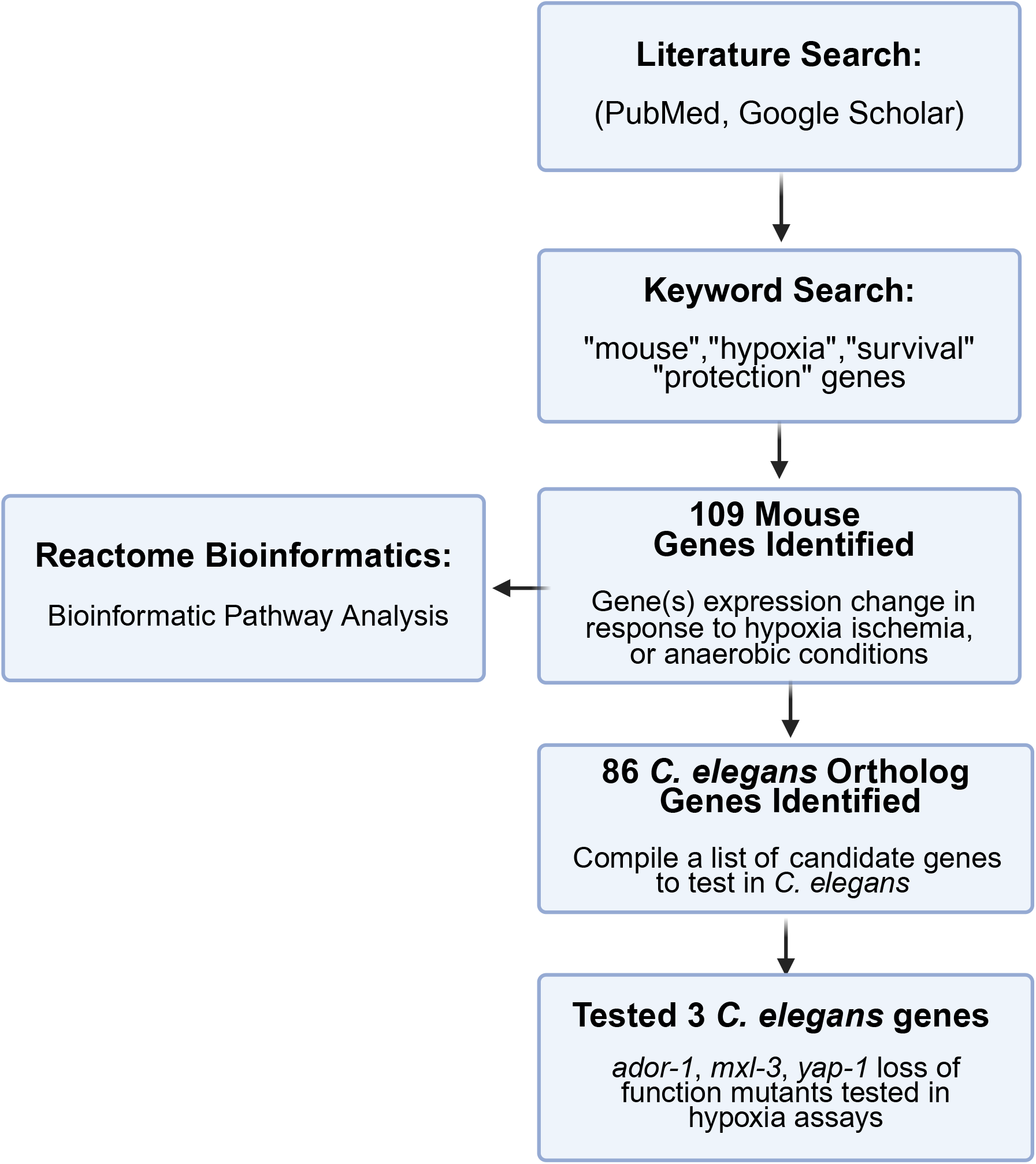
Schematic of The Workflow to Identify Candidate Genes. An overview of workflow used to identify mouse candidate genes and *C. elegans* orthologs.

The data was compiled and we identified human and *C. elegans* orthologs for each candidate gene using either Genome Alliance or DRSC Integrative Ortholog Prediction Tool (DIOPT) using gene alignment scores, and bioinformatics prediction tools that compare the nucleotide sequences across species to determine similarity and conservation of gene sequence. In cases where multiple genes are listed in Genome Alliance or DIOPT, we list the ortholog gene with the best alignment match or highest DIOPT score. Alternatively, where we list more than one ortholog gene, we found that these genes all had the same alignment score. In all other cases, only one ortholog gene is listed. A complete list of candidate genes and orthologs is provided as supplemental materials (Supplemental Tables 1).

We found at least 109 genes that were transcriptionally changed in response to hypoxic and ischemic conditions (Table 1). We used gene ontology databases and Reactome pathway analysis to determine the most prevalent cell pathways associated with the genes identified in our meta-analysis (Griss et al. 2020; Ragueneau et al. 2026). Based on Reactome analyses, these individual cellular pathways could broadly be grouped into cell pathways associated with (1) immune regulation, (2) general signal transduction, (3) gene expression changes in pathways that regulate metabolism and in disease states, and (4) development (Supplemental Table 2). Many of these pathways have been observed in mammalian hypoxic and ischemic cell states (Firth et al. 1994; Majmundar et al. 2010; Shay and Celeste Simon 2012; Lee et al. 2020; Luo et al. 2022). Our ability to identify genes like Hif1, PDK1, LDHA, and BNIP3L, all of which have identified roles in mounting a response to or provide protection to hypoxic ischemic insult (Semenza 1996; Semenza 2001; Kim et al. 2006; Papandreou et al. 2006; Bellot et al. 2009) provides confidence that our approach successfully identifies genes involved in regulating response to hypoxia and ischemia. We assembled a list of genes that regulated hypoxia and anaerobic conditions in mice, see Table 1, and noticed that there were some that had not yet been tested in *C. elegans*. We reasoned that if the candidate genes had a conserved role in adapting to hypoxia, then we would observe hypoxia-specific phenotypes in *C. elegans* with mutations in these genes. We prioritized which *C. elegan*s mutants to test in our assays based on the following criteria: (1) a genome alignment or DIOPT score of 4 or higher to increase the likelihood that we had identified a true ortholog, (2) candidate genes in contention associated with cellular stress responses, regulation of development, immunity, and/or metabolism. These are the most significant pathways found in our analysis, and (3) no previously reported phenotypes in hypoxia or anoxia. The three candidate genes we chose were *mxl-3*, *ador-1*, and *yap-1*.

**Table 1:**
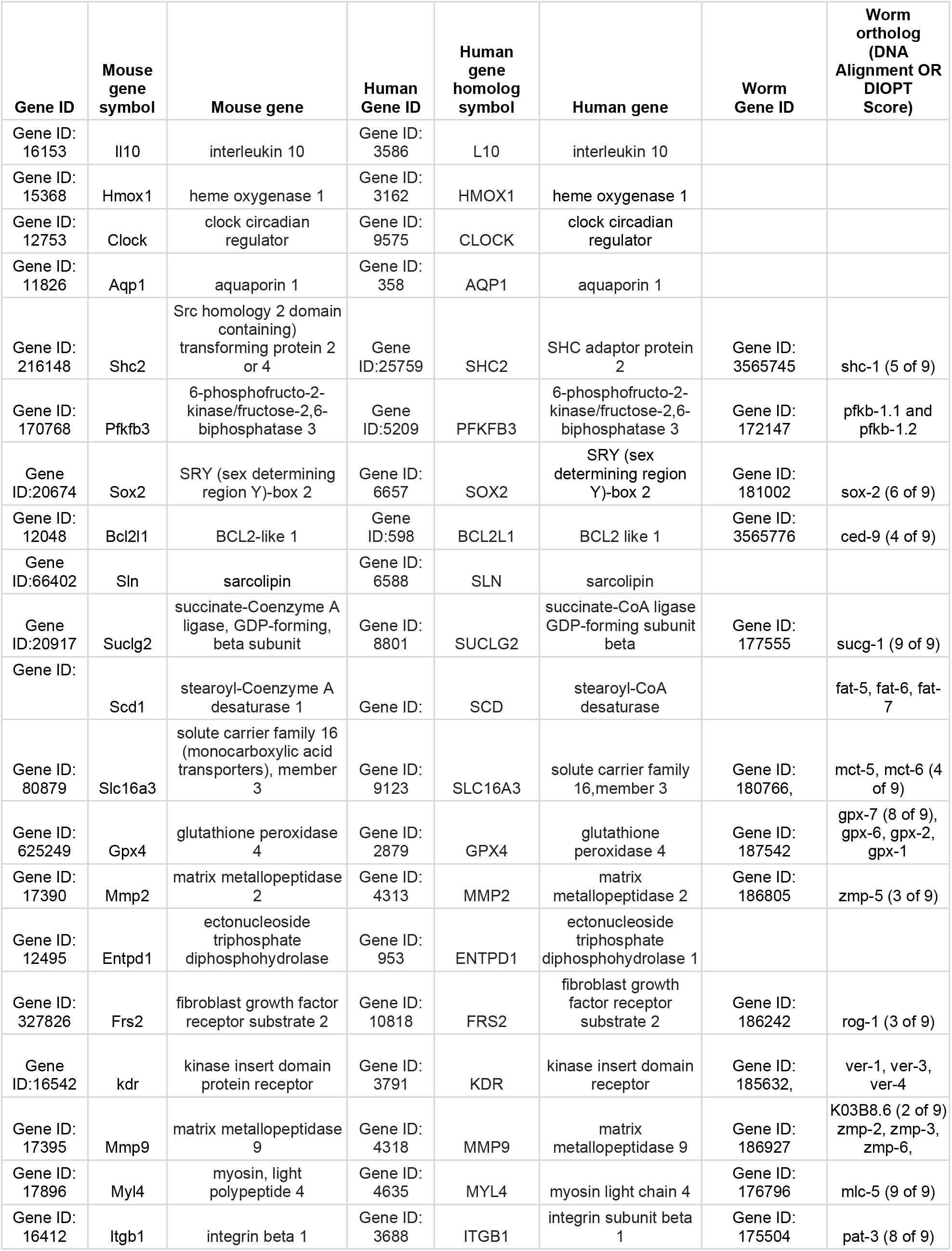
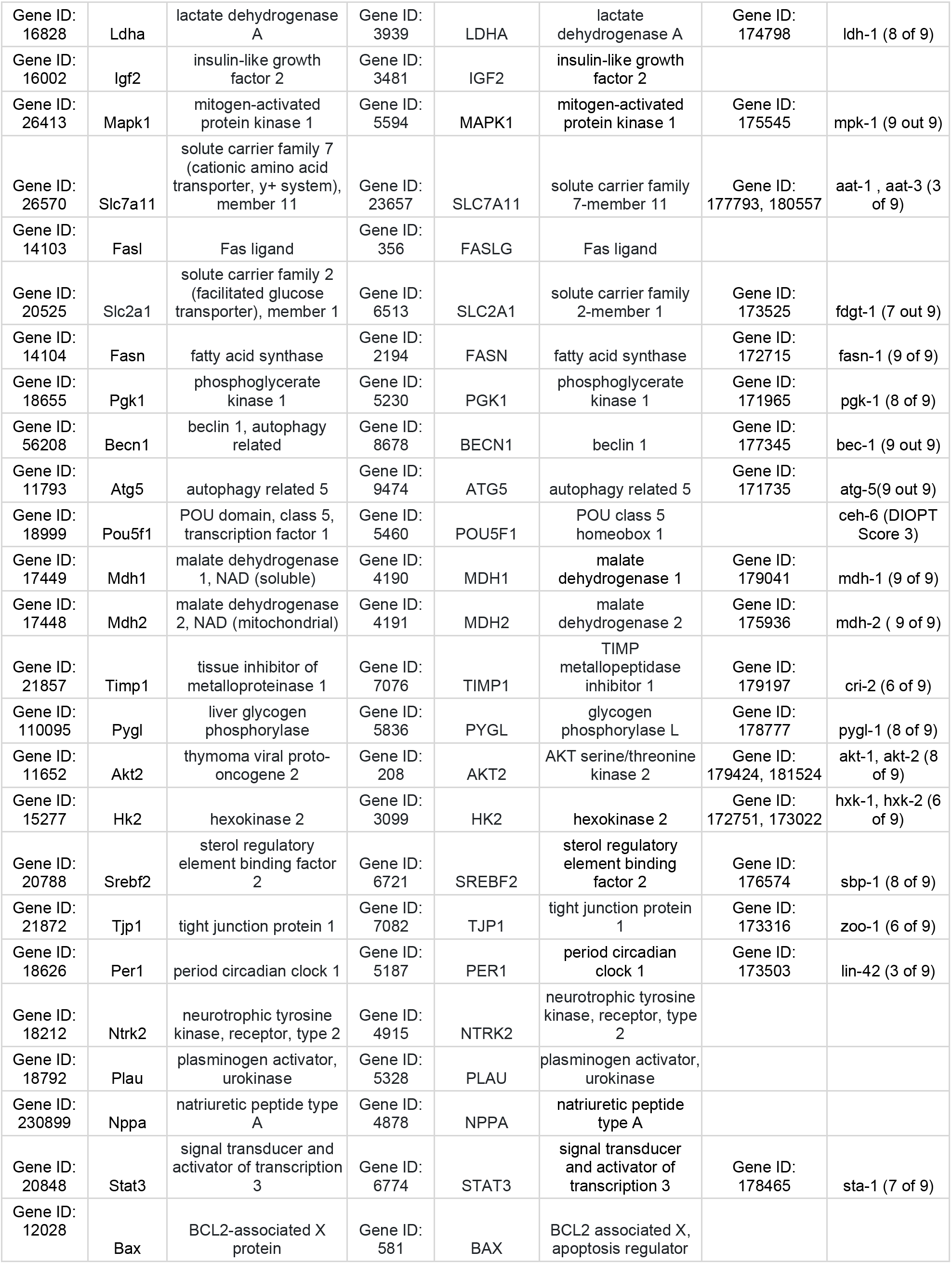
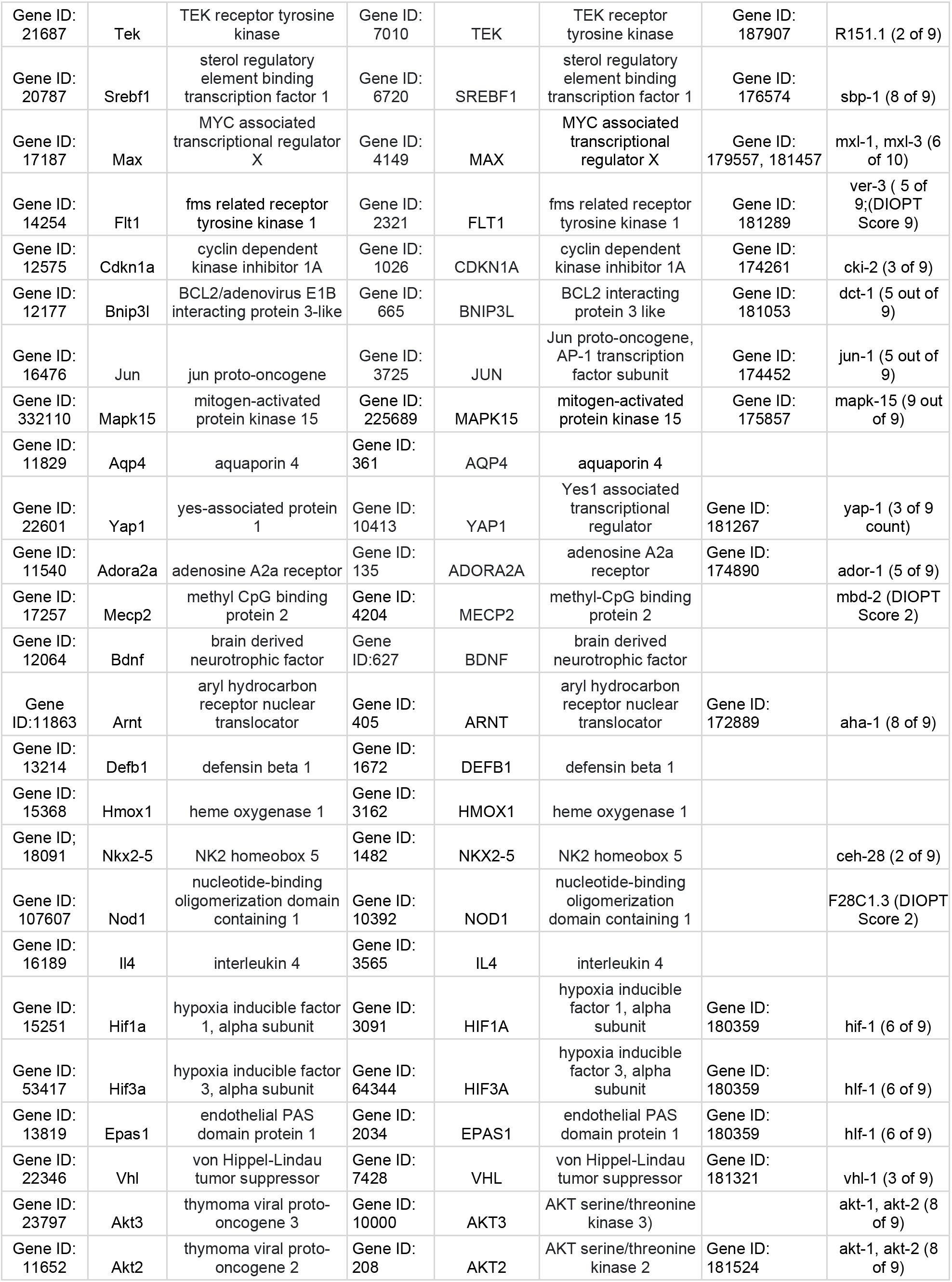
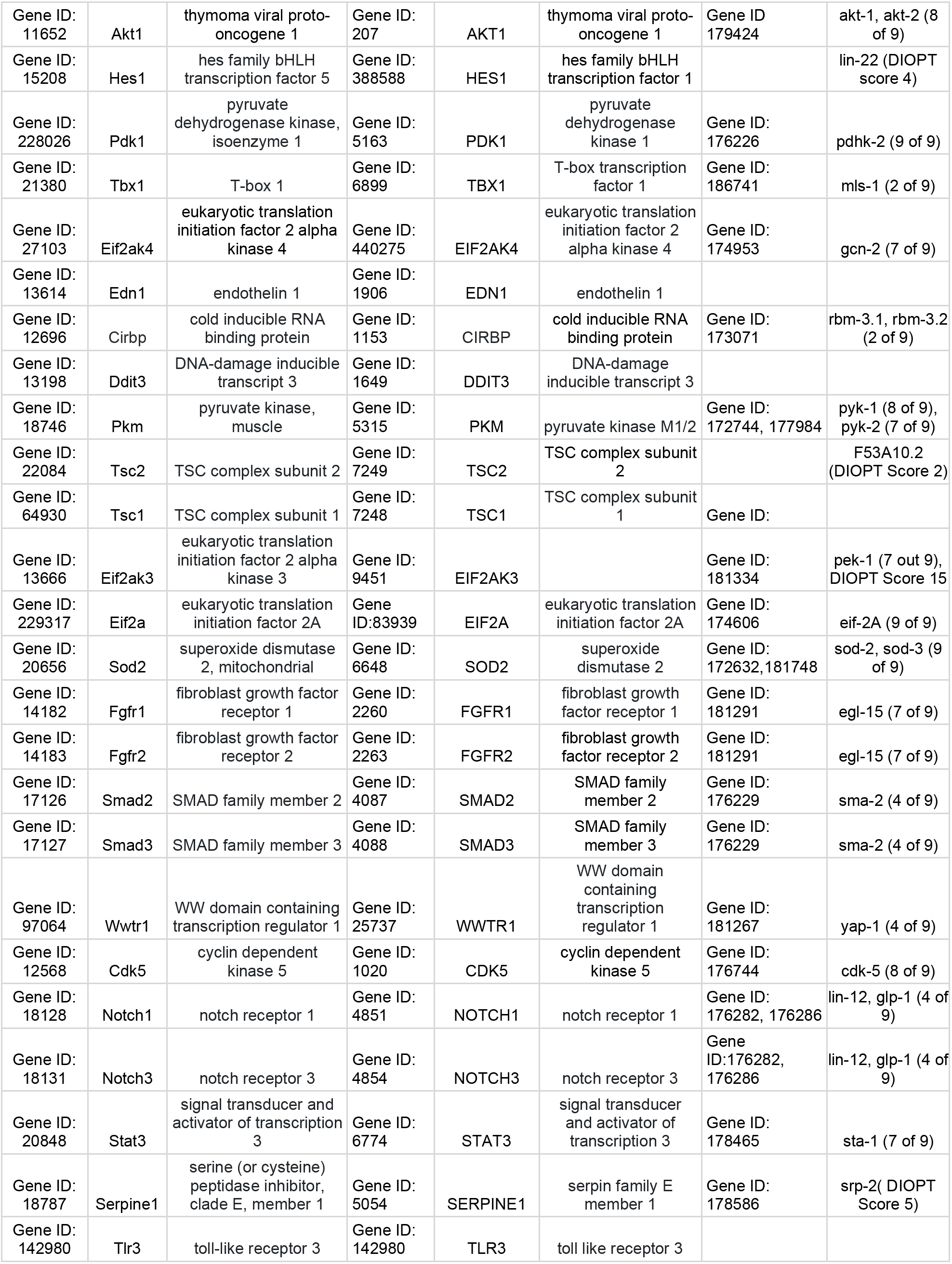
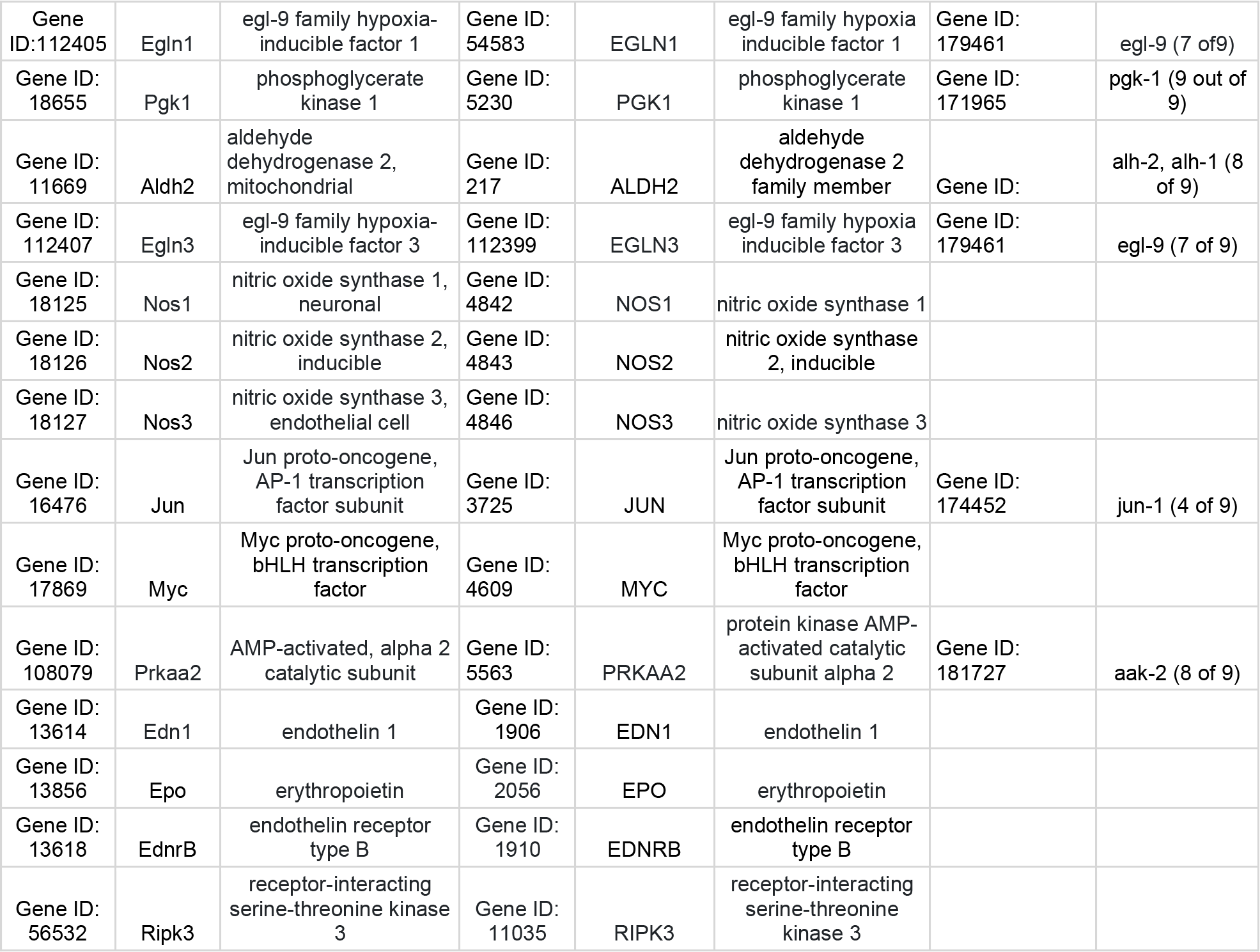
List of Mouse Genes Involved in Adaptation or Protection to Hypoxic Conditions. Human homologous and *C. elegans orthologous* candidate genes listed with NCBI Gene ID identifier.

*mxl-3*, a Myc-associated factor X (MAX) transcription factor belongs to a family of transcription factors that contain basic helix-loop-helix motifs (b-HLH) and leucine zipper domains (Lüscher and Vervoorts 2012). *mxl-3* regulates lipid metabolism, nutrient availability, and fat storage (Cuervo 2013). *ador-1*, a G-protein coupled receptor (GPCRs), and the only known adenosine receptor in *C. elegans*. Adenosine signaling through *ador-1* provides protection to oxidative stress (Ling et al. 2022), and there is evidence to support genetic interactions of common regulators of oxidative and hypoxic stress responses (Scott et al. 2002; Lee et al. 2010; Feng et al. 2021). *yap-1*, the ortholog to the human Yes-associated protein (YAP) (Huang et al. 2005; Lei et al. 2008) and functions downstream in the Hippo signaling pathway. YAP-1 is required for regulating and maintaining developmental processes, specifically cell polarity to establish neuronal polarization along the anterior-posterior (A-P) axis (Lee et al. 2018). In addition to roles in development, *yap-1* gene function is essential for *C. elegans* to mount effective immune and thermal stress responses (Iwasa et al. 2013; Ma et al. 2020).

We acquired *C. elegans* strains with loss-of-function mutations in *ador-1, mxl-3*, and *yap-1* so we could assess the role of each gene in hypoxia. We chose to assess the function of these genes in anoxia, 0.1% O_2_, and 0.5% O_2_ because these conditions activate physiologically and genetically distinct responses (Nystul and Roth 2004; Miller and Roth 2009; Iranon and Miller 2012). We tested both first-stage larvae (L1) and fourth-stage larvae (L4), because previous work has shown that sensitivity to oxygen deprivation varies with developmental stage (Padilla et al. 2002).

We first assessed whether these mutants were able to enter and exit from suspended animation. We assessed viability to gravid adults after a 24h exposure to anoxia as either L1 or L4. We did not observe a defect in any of the mutants we tested (Figure 2a, 2b), indicating that these genes are not required for the response to anoxia. We used the same test to assess viability after exposure to 0.1% O_2_, conditions that induce a reversible post-embryonic developmental arrest (but not suspended animation) in wild-type *C. elegans* (Miller and Roth 2009). All of the mutants tested tolerated 24h exposure to 1000 ppm O_2_ as well as wild-type controls (Figures 2c-d). These results indicate that *ador-1*, *mxl-3*, and *yap-1* are not required to survive severe oxygen deprivation.

**Figure 2:**
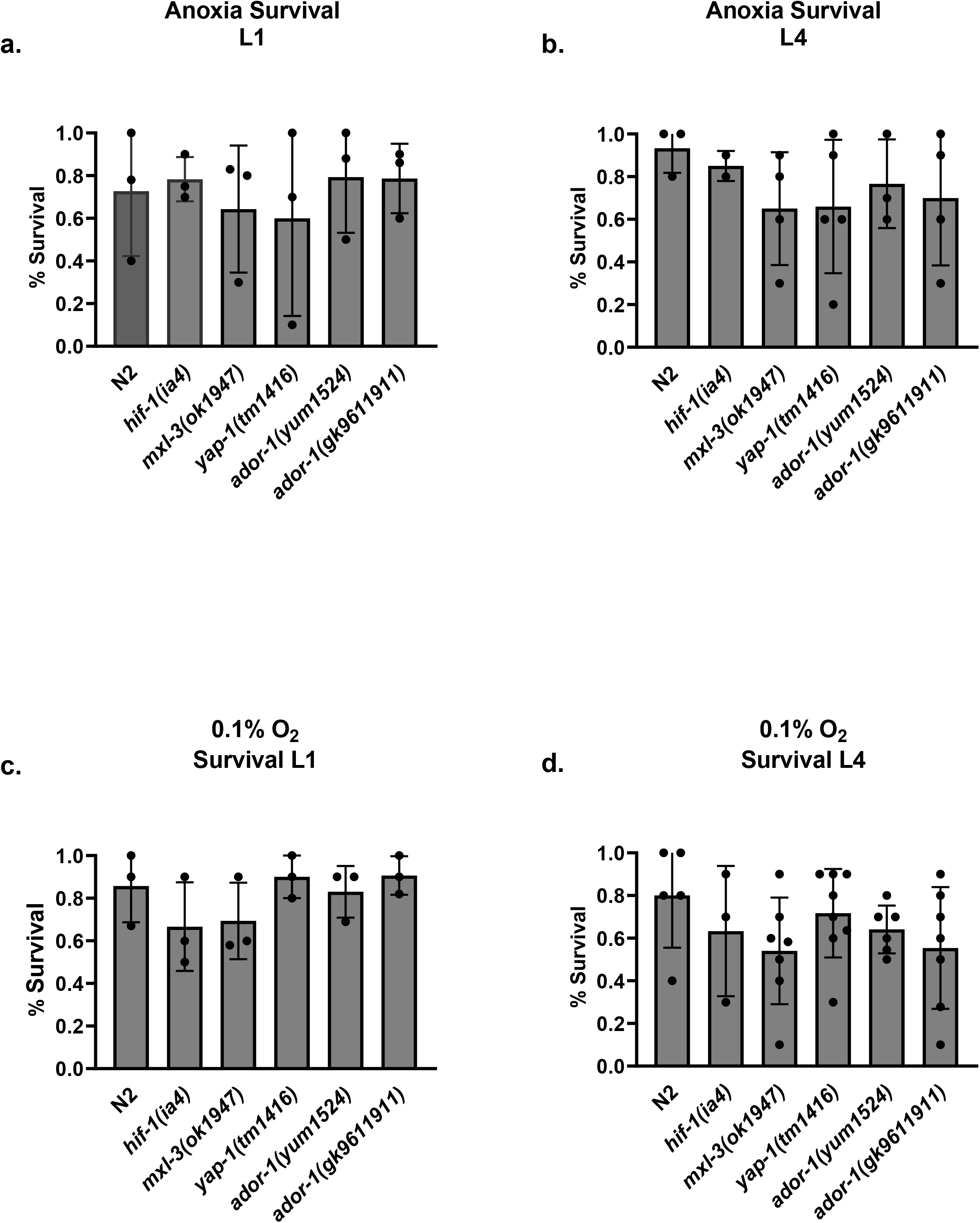
Loss of *mxl-3*, *ador-1*, and *yap-1* Gene Function Does Not Perturb Survival to Acute Hypoxia. **A-B.) Survival to Anoxia is Not Affected by Loss of either *mxl-3*, *yap-1*, or *ador-1*.** Synchronous (a) L1s and (b) L4s were obtained from adults (N2, *hif-1(ia4lf)*, *mxl-1(ok1947lf)*, *ador-1(yum1524 null)*, and *yap-1(tm416null)*) were allowed to lay eggs for 2-6 hours and the embryos were allowed to hatch at room temperature for 16-20h before hypoxic exposure. L1s were exposed to <10 ppm O_2_ for 22-27h at room temperature. After removal from hypoxic conditions, animals were allowed to recover at room air, room temperature, then evaluated 48-60h for the number of animals that developed into gravid adults. Results are shown for 3 independent experiments, data points show mean survival for 10 to 15 animals for each genotype. The bars represent the average mean for three experiments. No statistical significance assessed by one-way Anova followed by Tukey post hoc analysis; error bars represent the standard deviation of the mean. **C-D.) Loss of *mxl-3, ador-1,* and *yap-1* Does Not Impair Survival to 0.1% Hypoxia.** (c) L1s and (d) L4 animals were exposed to 1000 ppm O_2_ for 22-27h at room temperature and survival was scored after recovering in room air for 24-36h.Survival was assessed for the number of animals that survived and how many progressed to normal-looking gravid adults. Results shown for 3-5 experiments, data points show mean survival for each genotype. No statistical significance assessed by one-way Anova followed by a Tukey post hoc analysis; error bars represent the standard deviation of the mean.

In 0.5% O_2_, *C. elegans* continues developmental and reproductive activity, which is quite different from the reversible arrest of activities in more severe oxygen deprivation (Nystul and Roth 2004; Miller and Roth 2009). We found that L1 and L4 survived 24h exposure to 0.5% O_2_ as well as wild-type controls (Figures 3a-b), indicating that these genes are not required for the acute response to these hypoxic conditions. We also asked if these mutants were able to maintain continued reproductive activity in 0.5% O_2_. In these conditions, wild-type animals continue to fertilize oocytes and lay eggs but these processes are arrested in *hif-1(ia04)* animals (Miller and Roth 2009). We found that the rate of egg-laying in 0.5% O_2_ was similar for all mutant strains (Figure 3c). Moreover, all of the mutant animals were able to resume normal rate of egg-laying when returned to normoxia (Figure 3c). These results suggest that *ador-1*, *mxl-3*, and *yap-1* are not necessary to adapt reproductive activity during acute exposure to moderate hypoxia.

**Figure 3:**
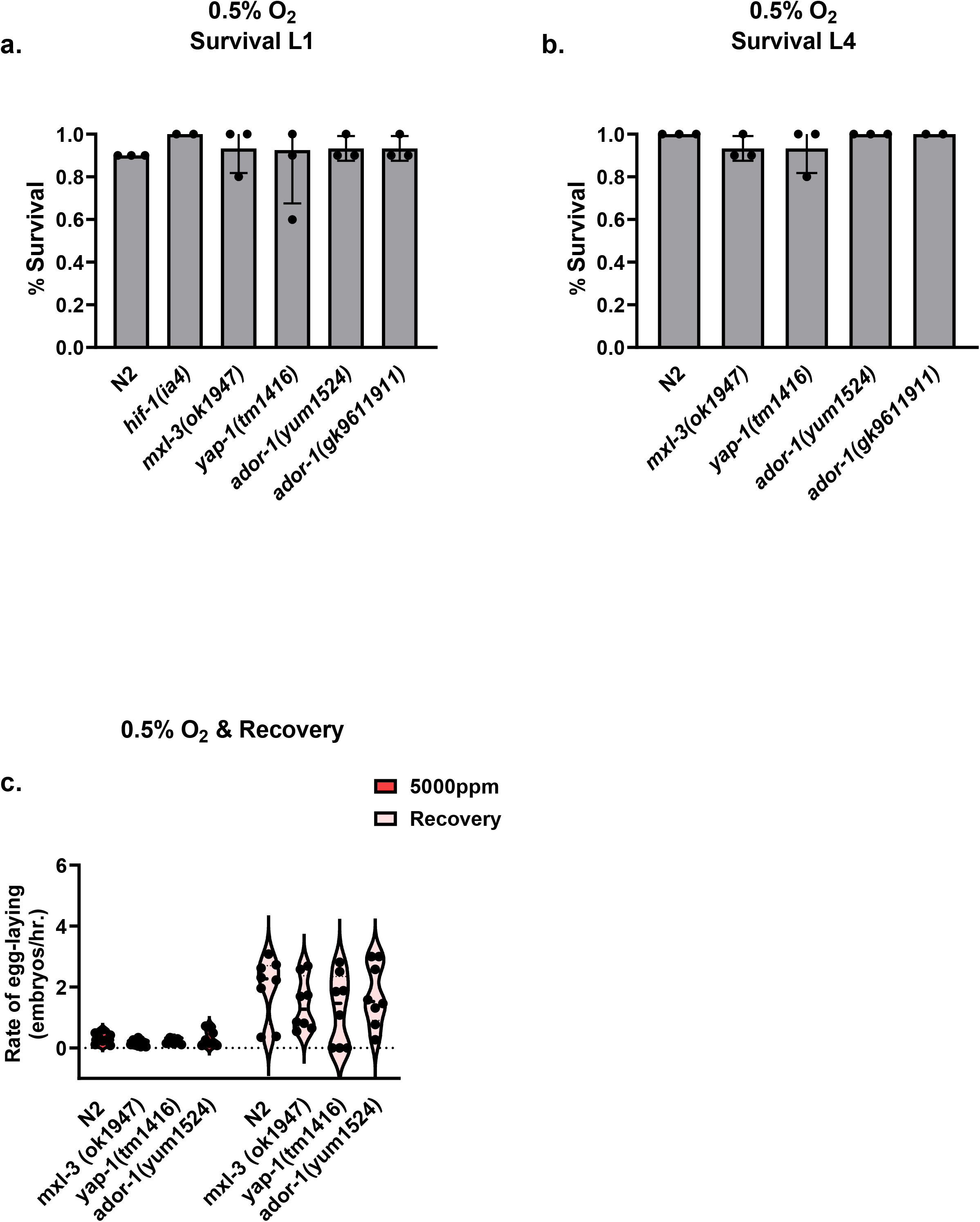
Neither Survival or Egg laying is Not Disrupted During 0.5% Hypoxic Insult. **A-B.)** (a) L1s and (b) L4 animals were exposed to 5000 ppm O_2_ for 22-27h at room temperature and survival was scored after recovering in room air for 24-36h.Survival was assessed for the number of animals that survived and how many progressed to normal-looking gravid adults. No statistical significance assessed by one-way Anova followed by a Tukey post hoc analysis; error bars represent the standard deviation of the mean. **C.) Rate of egg laying Is Not Altered In 0.5% Oxygen.** The rate at which 1 day adult wild type, as well as *mxl-1(ok1947lf)*, *ador-1(yum1524 null)*, and *yap-1(tm416null)* mutant animals lay eggs at 5000 ppm O_2_ for 24 hours. Animals were allowed to recover in room air for 24 hours and the number of eggs was counted. 10-15 animals are exposed to 5000 ppm O_2_, on 8 plates per genotype. Rate of egg laying is calculated for individual animals and the mean is calculated for each plate. Data shown for mean for each plate across genotypes, for 5000 ppm O_2_ and room air recovery. No statistical significance assessed by two-way Anova followed by a Bonferroni post hoc analysis in either condition.

We reasoned that since the effect of hypoxia can depend on the duration of exposure, we were motivated to assess the effect of longer exposure to moderate hypoxia. To assess the response to chronic hypoxia, we exposed L4 to 0.5% for up to 90 hours and then measured the total number of viable progeny the animals produced. In this assay, the L4 animals develop to adults: they complete vulval development, transition from spermiogenesis to oogenesis, and begin ovulation, fertilization, and egg-laying during the hypoxic exposure. Importantly, there is no significant difference in brood size associated with mutations in *ador-1*, *mxl-3,* or *yap-1* when animals remained in normoxia (Figure 4a).

**Figure 4:**
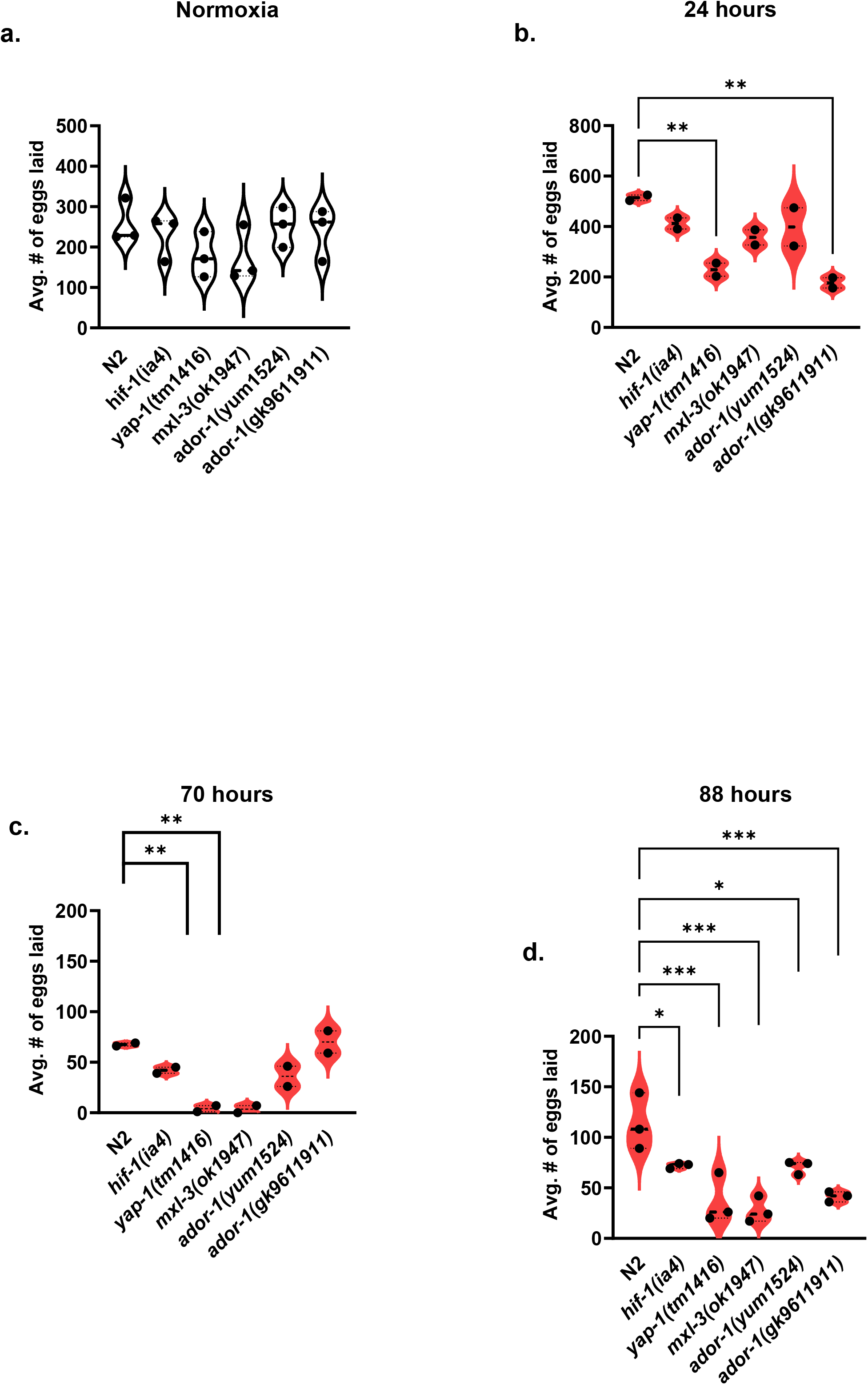
Loss of *mxl-3* and *yap-1* Perturbs Egg Laying Abilities with Prolonged Hypoxic Exposure. **A.) The number of eggs is not perturbed in L4 control animals.** Number of eggs laid in *hif-1*, *mxl-3, ador-1,* and *yap-1* mutant animals when left to develop into adults under normoxic conditions. Results shown for 3 experiments, no significance assessed by one-way Anova with a Bonferroni post hoc analysis. **B.) Exposure to 5000 ppm (0.5%) O_2_ in L4 y*ap-1* loss of function mutant animals results in a decrease in the number of eggs laid post hypoxia.** N2 wild type, *hif-1, mxl-1*, *ador-1*, and *yap-1* were picked as L4 animals and exposed to 5000 ppm O_2_ for 22-27 hours and allowed to recover in room air for 24-36h and scored for the number of animals that developed into normal-looking gravid adults. Animals were allowed to lay eggs for 2 to 6 hours. Results are shown for two independent experiments, the dashed line represents the mean. * Denotes significance assessed by one-way Anova followed by a Bonferroni post hoc correction. P value, < 0.001**. C-D) Prolonged Exposure to Hypoxia Further Decreases Egg-laying in *mxl-3* and *yap-1* Mutant Animals.** Experimental setup as described in panel C. Animals exposed to 5000 ppm O_2_ for 70 hours (c) or 88 hours (d) and assessed for egg laying post hypoxia. * Denotes significance assessed by one-way Anova followed by a Bonferroni post hoc correction. * P value, < 0.001, *** p value versus N2 wild type.

We observed a significant decrease in the viable progeny produced by *yap-1* mutants after a 24h exposure (Figure 4b), and this defect was exacerbated by longer hypoxic exposure (Figure 4c-d). We also found that *mxl-3* mutant animals were less able than controls to maintain progeny production after chronic hypoxia (Figure 4c-d), although this was not apparent after only 24h exposure. We also observed a hypoxia-associated decrease in progeny produced by *ador-1(gk9611911)* mutant animals after 24h exposure; however, this phenotype was not observed with another allele, *ador-1(yum1524*). The source of this discrepancy is unclear. Both of these alleles of *ador-1* are putative null alleles (Thompson et al. 2013; Pu et al. 2023). It may be that these mutants have a more variable response to hypoxia; neither mutant was statistically different from wild-type controls after 70h exposure to 0.5% O_2_, but both were affected after 88h (Figure 4c-d).

## Discussion

Here we describe a meta-analysis approach, where we survey the published literature for mammalian genes with focus on mouse studies, to identify genes that protect against the deleterious effects of hypoxic ischemic insult. Using online bioinformatics analysis and genome alignment software we have identified cross species (mouse, human, and *C. elegans*) candidate genes and cellular pathways that confer protection to hypoxic ischemic insult. Several mammalian studies investigating hypoxic and ischemic insults have broadly implicated signal transduction pathways dedicated to responding to diverse types of cellular stress, gene expression changes within metabolic pathways, as well as those that regulate cell cycle and development (Luo et al. 2022), and we find the same pathways and categories associated with our candidate genes.

This more targeted approach has a higher throughput and increases the possibility of finding genes whose perturbations provide a phenotype in our assays. We identified 86 *C. elegans* candidate genes, 51 genes of which have not been tested for a phenotype to hypoxia. We screened complete loss of function alleles in three *C. elegans* genes: *ador-1*, *mxl-3*, and *yap-1* under acute, moderate, or chronic hypoxic conditions. We find *ador-1*, *mxl-3*, or *yap-1* function is not required to survive anoxia, nor is gene function essential for animals to survive acute or moderate hypoxia. Unlike *hif-1* mutants, which suspend reproductive processes in moderate hypoxia (0.5% O_2_), *ador-1*, *mxl-3*, or *yap-1* mutant animals maintain reproductive activities. However, we show *yap-1* and *mxl-3* loss of function mutants reproductive abilities were compromised, and these animals produced less viable offspring, when exposed to chronic hypoxia, *i.e.,* longer exposure to moderate hypoxic conditions. Future studies will investigate how *mxl-3* and *yap-1* regulate reproductive processes, and how these cellular and molecular mechanisms are perturbed under hypoxic conditions. Additionally, our results highlight the differences in response and vulnerability in tissues with changes in acute versus chronic hypoxic states. Similarly, in cardiovascular diseases, the duration and severity of oxygen deprivation, leads to differing responses and susceptibility amongst tissues (Lucero García Rojas et al. 2021).

The molecular mechanisms that drive responses in acute and chronic hypoxic states are multifaceted and are difficult to fully investigate in more complex genetic systems. Our approach highlights a method to study hypoxia dependent phenotypes and investigate the underlying mechanisms in the genetically tractable nematode *C. elegans*.

## Data Availability

All strains are available at the Caenorhabditis Genetics Center. All raw data can be found in the supplement or available by request.

## Acknowledgements

We thank the Caenorhabditis elegans Genetics Stock Center (CGC), which is funded by the National Institutes of Health (NIH) Office of Research Infrastructure Programs (P40 OD010440). Some strains were provided by the Japanese National Bioresource Project. We would like to thank members of the Bennett Lab, Emily Jasmin, Marcellino Rau Nikhil Desai for providing intellectual support in our literature review analysis. We would like to thank Izabel Kickner, alumnus of Bard College, for providing ideas at the beginning of the project. We would like to thank Anne C. Hart for providing intellectual support and feedback in our results.

H.L.B and D.L.M designed the experiments. E.M., G.Z.W., K.N., D.L.M, and H.L.B conducted experiments and/or constructed figures. H.L.B analyzed all results and H.L.B and D.L.M wrote the manuscript. All authors proofread manuscript.

